# Testing the waters of macrophyte biodiversity with multiscale spatial analysis of public lake monitoring data

**DOI:** 10.64898/2026.06.22.733670

**Authors:** Misha Tseitlin, Jorge García-Girón, Julie Crabot, Xiaoming Jiang, Daniel J. Larkin, Janne Alahuhta

## Abstract

Freshwater monitoring programmes like the European Union’s Water Framework Directive (WFD) provide a wealth of data on European lake status, including water quality and macrophytes (aquatic plants) as critical habitat features that support health of humans and wildlife. Easier WFD data access can enable external management and research to better safeguard human and natural freshwater use. We demonstrate a replicable workflow to easily download and process multi-year (2007–2024) observations of lake macrophytes (425 sites) and complementary water quality variables (202 sites) from Swedish WFD data. Then, we illustrate the value of improved data access to address ecological questions that drive conservation, investigating how spatial scales influence macrophyte richness and associated water quality relationships using a spatial random intercept model. Decomposing the spatial intercept links small scales (<10 km) to site-level gradients and large scales (>100 km) to biogeographical drivers. Stochastic and environmentally-structured processes coexisted at intermediate scales (10–100 km). Adding water quality rarely improved overall predictive performance of macrophyte diversity models but consistently influences the role of different spatial scales. Water quality variables showed consistent spatially structured variation at intermediate scales and unique spatial patterns in tandem, overlapping with large-scale biogeographical influences. Altogether, we show context-dependencies for spatial model interpretation and provide guidance in accounting for spatial confounding to improve inferential and predictive performance. Our workflow and results show a clear way forward for accessing high-quality macrophyte and water quality data sets and their utility for addressing ecological questions that guide macrophyte protection under the WFD.

**Highlights:**

- years Swedish of macrophyte and water quality monitoring data were extracted.
- richness showed scale-specific patterns linked to geographic gradients.
- best predictive models for richness had no water quality at all.
- overlap in their spatial scales and must be carefully separated.
- pen access data and multiscale analysis can apply to many ecological questions.

## 1. Introduction

Macrophytes are foundational species in freshwater aquatic ecosystems worldwide: their high biomass and abundance define shallow areas (i.e., 1.5–5 meters), provide resources to other organismal groups, and physically structure the environment (Bolpagni, 2026). Diverse macrophyte communities support high abundance of conservation-targeted species, as well as other fish, invertebrate, bird, and semi-aquatic mammal species through habitat formation (Bolpagni, 2026; Larson et al., 2020). Positive associations between plant biodiversity and water quality — partly due to biofeedback mechanisms through which macrophytes regulate and improve water quality — allow macrophytes to serve as general surrogates for ecological and hydrological conditions in freshwater systems (Law et al., 2019). These processes include nutrient cycling via uptake and sequestration, water purification, sediment stabilisation that reduces turbidity, oxygen production, and the modulation of biogeochemical feedbacks that collectively govern water quality. Because of their wide-ranging ecological roles, the EU’s Water Framework Directive (WFD) mandates macrophyte monitoring, which can inform and support nature-based solutions (NbS) to halt freshwater biodiversity collapse (Bolpagni, 2026).

WFD, Infrastructure for Spatial Information in Europe (INSPIRE), and other international monitoring programmes produce extensive macrophyte and limnological datasets with high value for capturing macrophyte biodiversity. However, traditional workflows for data retrieval, cleaning, and integration create bottlenecks in research progress. Despite EU reporting mandates, accessibility of freshwater system data lags behind that of other ecological indicators (Markovinović et al., 2022). When available, freshwater data exhibit nuances based on their local contexts, requiring complex preparation and assumption-laden, manual harmonisation. Application programming interfaces (APIs) offer an opportunity for data delivery with fast, reproducible pipelines and streamlined workflows for research and policy support. Of course, accessible data being in public repositories increases their use, especially for research (Pivari et al., 2019). Increasing accessibility of datasets and workflows would enable more ecological insights and use of existing long-term datasets that support policy, conservation, and management of freshwaters under the WFD.

WFD Swedish lake data capture large gradients of macrophyte richness, time, spatial extent, biogeography, and human influences (Ecke, 2018). Leveraging improved WFD data access and cleaning pipelines can allow for testing modern, non-parametric models that address critical macrophyte ecology gaps and the broader freshwater biodiversity crisis (Alahuhta et al., 2021a). Predictive models, such as spatially explicit non-parametric models, can capture heterogeneous and non-linear spatial structures that better approximate real-world ecological trends while minimising intrinsic and user-specified bias in conventional spatial models (Murakami et al., 2026a, 2026b). These models also support a classical linear structure, allowing incorporation of established ecological knowledge, comparison of multiple models, and evaluation of their assumptions about data structures to best predict macrophyte richness.

Macrophyte biodiversity operates across multiple spatial scales, including local (e.g., water depth, sediment type), lake (e.g., lake area, shoreline length), and biogeographical scales (e.g., catchment and latitude influences) (García-Girón et al., 2020). Thus, proper biodiversity prediction and monitoring rely on effective spatial scale identification — from standardised monitoring coverage to specific bioindicator selection (Gebler et al., 2024; González del Tánago et al., 2021). Here, we see an opportunity to clarify spatial relationships at national extents to improve monitoring efficiency and ecological understanding. Sweden presents an opportune testing ground. (1) Its relevance extends both to other EU WFD states but also across the Atlantic given previous research on cross-continental Nordic-US macrophyte diversity comparisons (Luukkonen et al., 2024). (2) Varied biogeographic conditions across topographic gradients and marked effects from Late-Quaternary glacial history across Sweden’s latitude pose potential for stronger generalisation (Lundqvist, 2004). (3) Strong spatial structures of land use in the Swedish central plains allow comparison to the European “agricultural heartland” while the north remains more analogous to Nordic forests (Winkler et al., 2025).

Water quality variables remain central to macrophyte macroecological analysis, but current findings largely remain correlative and associative (Correia et al., 2026; Iversen et al., 2019). Bidirectional causality (e.g., macrophyte-chemistry feedback loops) constrains inference despite broader methodological advances (Correia et al., 2026; Hilt et al., 2025). Conversely, predictive models offer practical value, but their performance may be scale-dependent, as disturbance regimes and environmental gradients often covary differently across spatial scales (Gebler et al., 2024). This phenomenon, where spatial imbalances appear even in sampling points (e.g., at site clusters at local scales), is often true for observational data in ecology (Diggle, 2013). Such confusion stems from spatial confounding, where model spatial components take explanatory power from linear ecological variables that function at overlapping spatial scales (Khan and Berrett, 2024). More simply, model over-reliance on aggregated spatial model components (e.g., basis functions, eigenvectors) can cause bias (Murakami et al., 2026a, 2026b; Ward and Anderson, 2026). These modelling shortfalls present an opportunity to leverage new spatial predictive methods, resolve existing tensions, and present a clear way forward to synthesise observational macrophyte and water quality data into a coherent modelling framework.

Our study uses WFD biomonitoring data from hundreds of Swedish lakes as a timely test case to share a repeatable workflow for improving data access and usability that is transferable to other similar ecological monitoring data. After preparing analysis-ready WFD Swedish lake data, we apply a novel multiscale model — CF-GLMM (coarse-to-fine generalised linear mixed model) — to address ecological questions regarding macrophyte biodiversity at multiple spatial scales across Sweden. We aim to (1) access, process, and scrutinise national-level WFD monitoring data for informing macrophyte ecology; (2) understand their utility to varied approaches for macroecological spatial modelling; and (3) decompose specific multiscale patterns and ecological drivers (i.e., nutrient status and water temperature) underlying Swedish macrophyte biodiversity and biogeographical trends. We subsequently abbreviate these as (1) data, (2) spatial, and (3) multiscale aims or objectives. These objectives create a repeatable workflow for other applications, identify the role of environmental (e.g., water quality) and biogeographic gradients relevant for macrophyte diversity. We further illustrate these gradients’ operational spatial scales and structures (i.e., often masked in single-scale models) in support of scalable monitoring and management decisions through the WFD. To that end, we discuss the multiscale method’s applicability to other macrophyte questions beyond our specific focus on Swedish lakes.

## 2. Methods

### 2.1. Data Workflow

To achieve the data aim, we document our workflow in detail and provide R scripts for easy data download and preparation in a companion GitHub repository (https://github.com/MishaTs/SwedenWFD). The workflow centres around Swedish freshwater macrophyte and water chemistry data from the Miljödata MVM system (https://miljodata.slu.se/MVM/). Users could either download data directly through the public search portal, which offers quick but limited CSV downloads, or download full, site level data via authenticated API access using a public token. We opted for the latter API approach. Our workflow retrieved, standardised, and cleaned the macrophyte dataset by removing internal quality fields, translating and harmonising variables, refining spatial and taxonomic information, and selecting only the first survey occurrences (i.e., for sites sampled multiple times) to reduce bias. We separately downloaded and processed water chemistry data, which was filtered to key parameters, unit harmonised, adjusted for detection limits (i.e., the lowest and highest chemistry values that can be determined based on the analytical protocols), and restricted to shallow sites with corresponding macrophyte records. Finally, we merged both datasets into an analysis-ready data product focussed on yearly macrophyte richness and associated water chemistry across hundreds of Swedish lakes. Our workflow made this unique data accessible and yielded a spatially comprehensive, multiyear dataset suitable for ecological assessments and macroecological models.

We successfully extracted data from 425 Swedish lakes from 2007 to 2025, of which half had some sort of water chemistry data (Table 1). Altogether, we provide an expanded dataset with indicators of macrophyte communities and chemistry that can be used to compare against and go beyond our subsequent analysis (Ecke, 2018). The coverage of key macrophyte and chemistry variables differed, as only a subset of monitored Swedish lakes have undergone structured macrophyte surveys since WFD implementation in 2000 (Fölster et al., 2014).

**Table 1:**
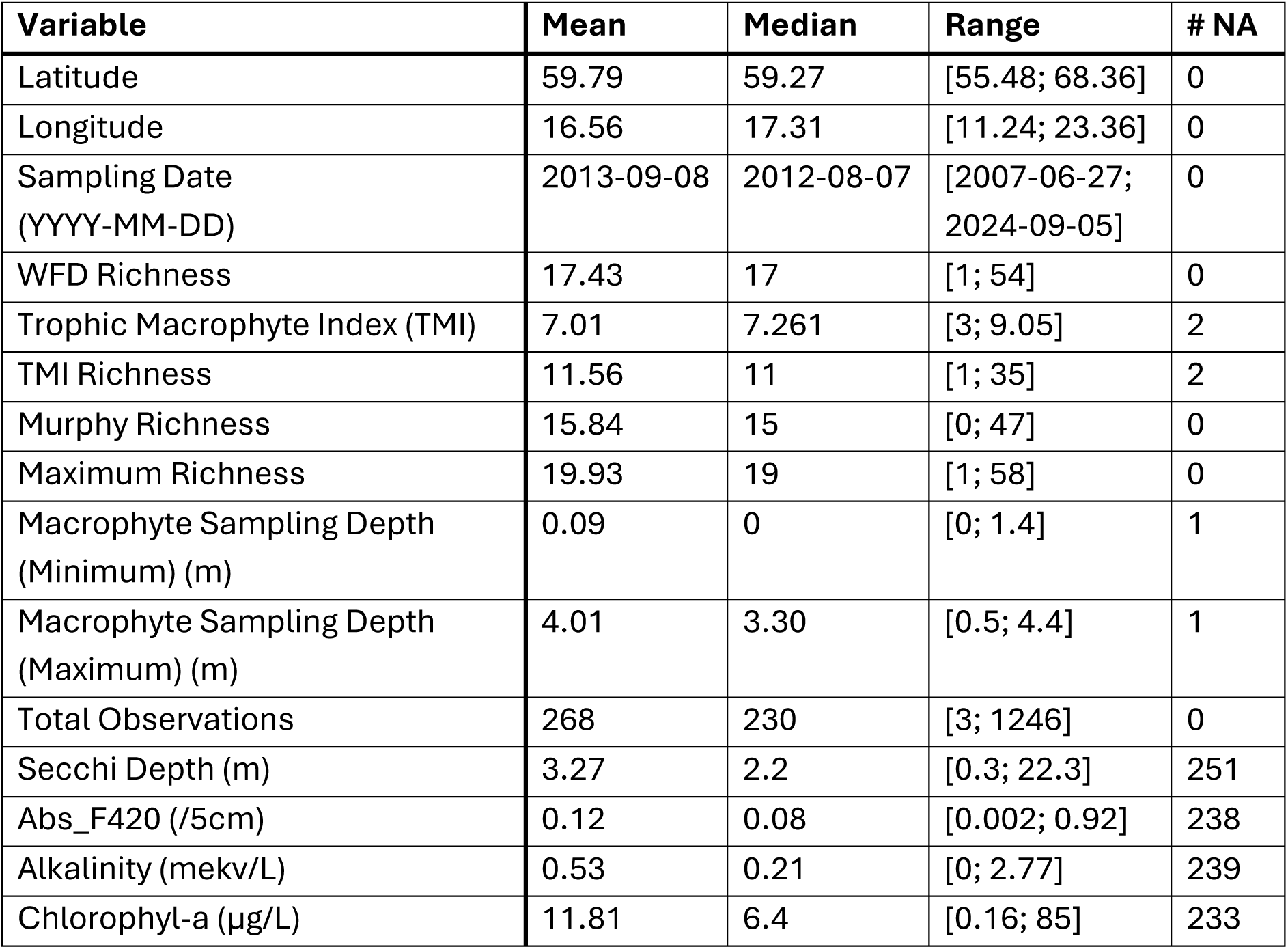
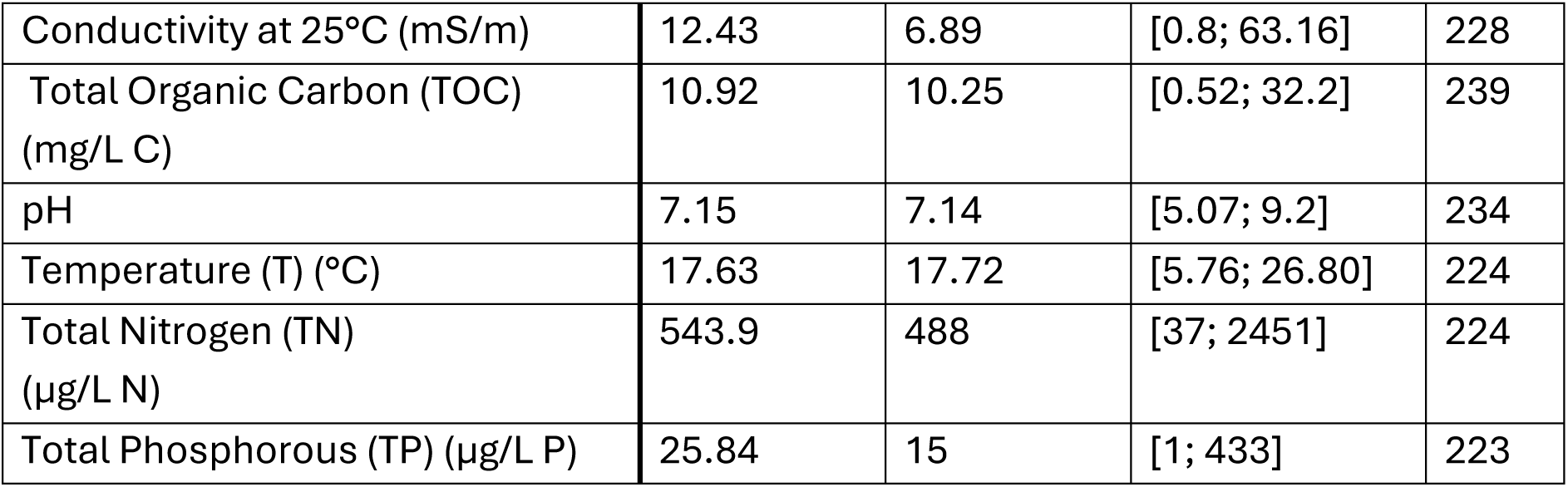
Summary statistics for main quantitative variables in the cleaned Swedish data used for analysis. Range presented in [min; max] format. Spatial coordinates in WGS84 (EPSG:4326). Data has 425 lakes total. Other available variables describe the sampling regime in more detail, including the survey protocol, sampling methods (i.e., rake, snorkel, binoculars, by-hand), and programme/study names.

Notably, chemistry data coverage linked strongly to the monitoring sub-programme. Looking at data with TN, temperature, and TP, only the NMÖ (National environmental monitoring) and KMÖ (Local environmental monitoring) programmes had chemistry for most observations. But the latter KMÖ only provided 10 sites, all in Stockholm County. RMÖ (Regional environmental monitoring) and KÖ (local monitoring) contributed the next greatest number of observations to the filtered data, respectively. Separately, we note that the SRK, RK, and KÖ programmes focus specifically on human-impacted water body monitoring with naming distinctions based on the surveyor(s), so they must be analysed in tandem with metrics of human activity (*Sveriges lantbruksuniversitet*, 2026).

### 2.2. Model Specification

To predict and decompose species richness patterns across multiple spatial scales (i.e., for spatial and multiscale aims), we used a modified generalised linear mixed model (GLMM) with a Poisson distribution for count data. Specifically, a coarse-to-fine (CF) GLMM provided a nice way to account for multiscale spatial trends:

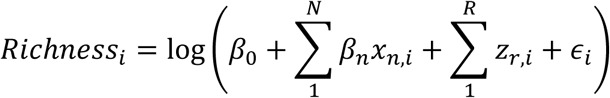

The main terms were a fixed intercept *β*_0_, coefficients for nth water quality variable *βn*, and coefficients for r^th^ local spatial scale *z_r_*, and error *ε*. The model estimated these for each site i, which corresponds to a row in our data. We compared models with chemistry coefficients against a null spatial model, which used just a fixed intercept (*β*_0_) along with each relevant spatial term (*zr*) to model richness. We define some of these key statistical terms with simple language in an appended glossary (Appendix A).

The CF mixed effects in our GLMM were the spatial functions **Z** (all *z_r_*), which together we call a spatial random intercept. Each distance r corresponded to some bandwidth (in metres). Rather than relying on user specification like traditional Moran eigenvector map (MEM), GAM, and stochastic partial differential equation (SPDE) models, holdout validation selected the optimal bandwidths corresponding to relevant spatial scales in the data. This helps CF models consistently outperform GLMs and MEMs. Under unknown spatial complexity, CF-GLMMs exceed GAM performance too (Murakami et al., 2026a, 2026b).

These models precisely estimated the functional spatial scales of Swedish macrophyte richness and their responses to adding and removing fixed effect terms (e.g., water quality). They did so while avoiding spurious fits that arise from user misspecification, a known problem for other decomposition-based methods and especially relevant for dispersed count data (Harrison, 2014; Veen, 2023). We used CF-GLMMs to explore macrophyte richness patterns at three scales: small (< 10 km), medium (10-100 km), and large (> 100 km). These scales contain roughly equal numbers of bandwidth components on a logarithmic scale and match the 3-scale breakdown from previous CF-GLMM analyses (Murakami et al., 2026a, 2026b).

### 2.3 Model Application

Before running the CF-GLMM, we selected the originally reported WFD richness and chemistry values during the macrophyte survey season. For chemistry, we calculated their weighted seasonal average and retained only water quality variables with less than 25% missing values.

We implemented a CF-GLMM with the bespoke spCF R package (Murakami et al., 2026c). We used the Poisson family, as quasi-Poisson implementation gave identical parameter estimates and overall results. This package provided relatively limited *a posteriori* checks, with no distributional parameters on mean, variance, or dispersion. But we computed raw residuals (i.e., observed - fitted) to assess goodness of fit. Out of the three predictive metrics (i.e., validation pseudo R^2^, RMSE, and MAE), we focus on pseudo R^2^ (henceforth just R^2^) for easier interpretation but provide the other scoring metrics from our chemistry models (Table S1).

The spCF package did not tolerate missing values, so our fixed effects analysis of water chemistry variables only included total nitrogen (TN), water temperature (T), and total phosphorous (TP). As all three variables may demonstrate non-linear relationships, we also considered adding a squared term (Alahuhta et al., 2020). We compared these covariates to a null model fit on the same site subset using a “dredging” workflow: we fitted all possible candidate models with TN, temperature, TP, and their squared terms (i.e., 27 total including the null model) and reported their results.

In general, CF-GLMMs can suit different purposes depending on research objectives. Different metrics matter depending on the research questions — inferential (e.g., quantifying causal effects of water chemistry on species richness), predictive (e.g., predicting species richness to a new area), and/or spatial (e.g., uncovering the spatial scales that chemistry-richness relationships function on) (Figure 1). Because of our dredging workflow, we examined these differences *a posteriori* by interpreting evaluative metrics for model suitability: coefficient-specific significance guided inference on βs, and pseudo-R^2^ indicated full-model predictive performance. CF-GLMMs automatically select relevant spatial distances—the presence/absence of which we interpreted for our spatial and multiscale objective.

**Figure 1:**
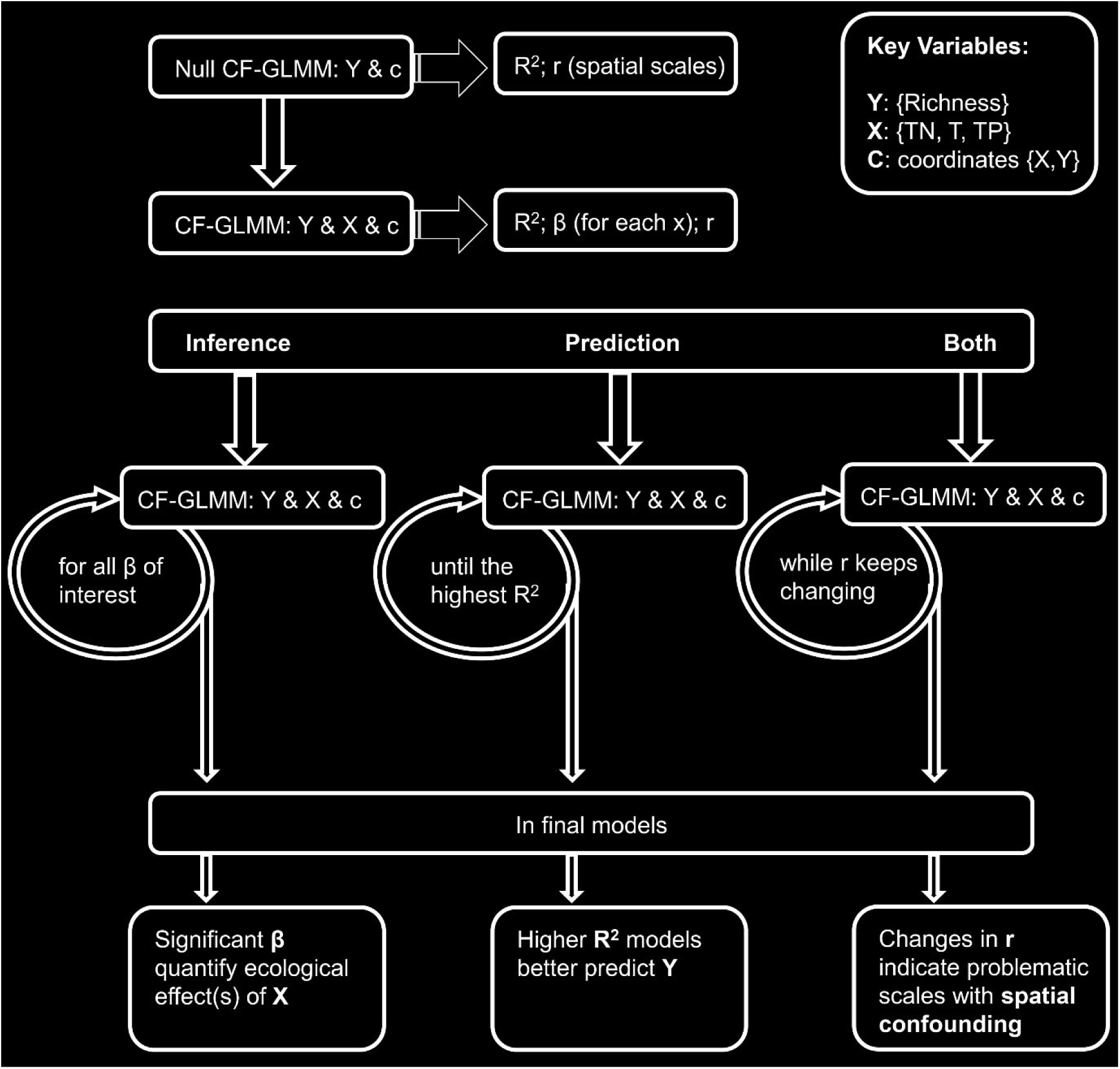
Conceptual representation of modelling workflow for CF-GLMMs highlighting differences depending on research objectives. As it can impact both prediction and inference, we address spatial confounding under the heading “both.” Terms include magnitudes β for all fixed effects, scales r corresponding to distance-specific bandwidths, and our X variables total nitrogen (TN), temperature (T), and total phosphorous (TP). We demonstrate the progression from a null model to a model incorporating fixed covariates X. Depending on the research objectives, the best metrics to evaluate will differ. We focus on X and Y variables from our analysis, but researchers can replace these with alternative variables as needed (e.g., pH and beta turnover, respectively). R^2^ represents general model adherence to the data between 1 (perfect fit) and 0 (no fit). Predictive scores like RMSE and MAE can provide an alternative to pseudo R^2^ for evaluating predictive performance. Main parameters are described in the upper right inset box.

## 3. Results

### 3.1. Data Coverage

Per our data objective, biodiversity and chemistry revealed strong spatial and macroecological patterns and yearly trends across the hundreds of Swedish lakes, affirming their utility for our spatial and multiscale analytical aims (Figure 2). Richness, TN, and T showed strong north-south and east-west gradients related to both sampling coverage and macroecological processes. The north had lower TN, T, and surveying density, and richness and nutrients increased slightly towards the northeastern coast. Sweden’s central lowlands host the country’s largest lake systems and population centres: combined with denser survey coverage and higher nutrient and temperature values, higher richness values often appeared in this region. Generally, the south-central east had both lower richness and TN compared to the south-central west. While showing some strong trends, variables remained heterogenous, especially outside the north, warranting more targeted analysis.

**Figure 2:**
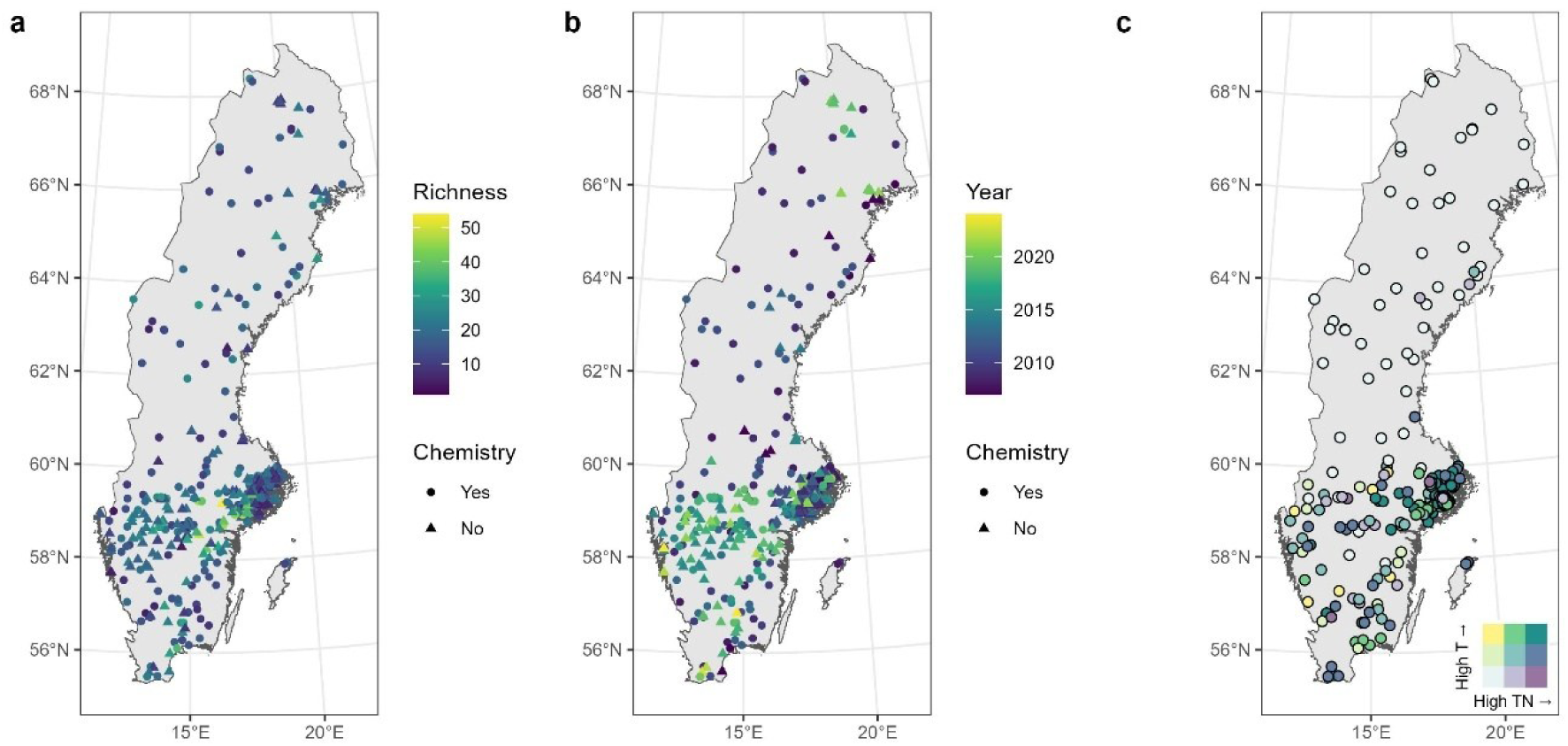
Spatial distribution of full and filtered data, highlighting key variables. a) Spatial distribution of species richness distinguishing between all monitored lakes, and those with suitable chemistry data, b) Spatiotemporal distribution of survey points, again distinguishing between data types. c) Bivariate plot showing 3 quantiles of temperature (T) and total nitrogen (TN) values and their co-occurrence across Sweden for only the non-missing filtered data subset. Legend in bottom right of the plot.

Some variables suggested structural spatiotemporal survey bias: eastern monitoring points were more likely to be monitored during later months, while points more north and east had their first surveys in earlier years (Figure S1). Richness also strongly correlated with the number of observations at a site. Macrophyte metrics were largely fit for purpose — WFD, vascular plant, and maximum richness cohered very strongly with minor differences. Notably, Sweden included some non-Murphy list species in its WFD richness metrics, and a few sites even had 0 Murphy richness (Table 1). Alkalinity, conductivity, and pH strongly correlated among themselves. TN and TP also related strongly, and TN had more general correlation with all water chemistry parameters.

### 3.2. Multiscale Richness Patterns

For the spatial objective, the CF-GLMMs illustrated the relevance of the three spatial scales driving macrophyte richness patterns (Figure 3). When combining the many spatial patterns into three macro-scales of small (< 10 km), medium (10-100 km), and large (> 100 km), distinct trends emerged for each. Small scales captured the variability (i.e., towards lower richness) around the major lake systems in Stockholm (59° N 18° E) and Vättern (58° N 14° E): these might correspond to spatial correlation from site proximity within single lakes or lake systems. Medium-scale patterns also trended towards lower richness, concentrated especially around urban Stockholm with gradual increases in richness away from the metropolitan area. Otherwise, the medium scales appeared to capture substantial stochasticity that we could not clearly link to any primary geographical patterns. In contrast, the large scales corresponded with dominant biogeographical trends, like higher richness along the coast and in the south-central valley and lower richness in the westward mountains and colder northern lakes. Together, the small and medium scales might reflect surveying biases and inherent statistical randomness in contrast to the clear, stable large-scale trends).

**Figure 3:**
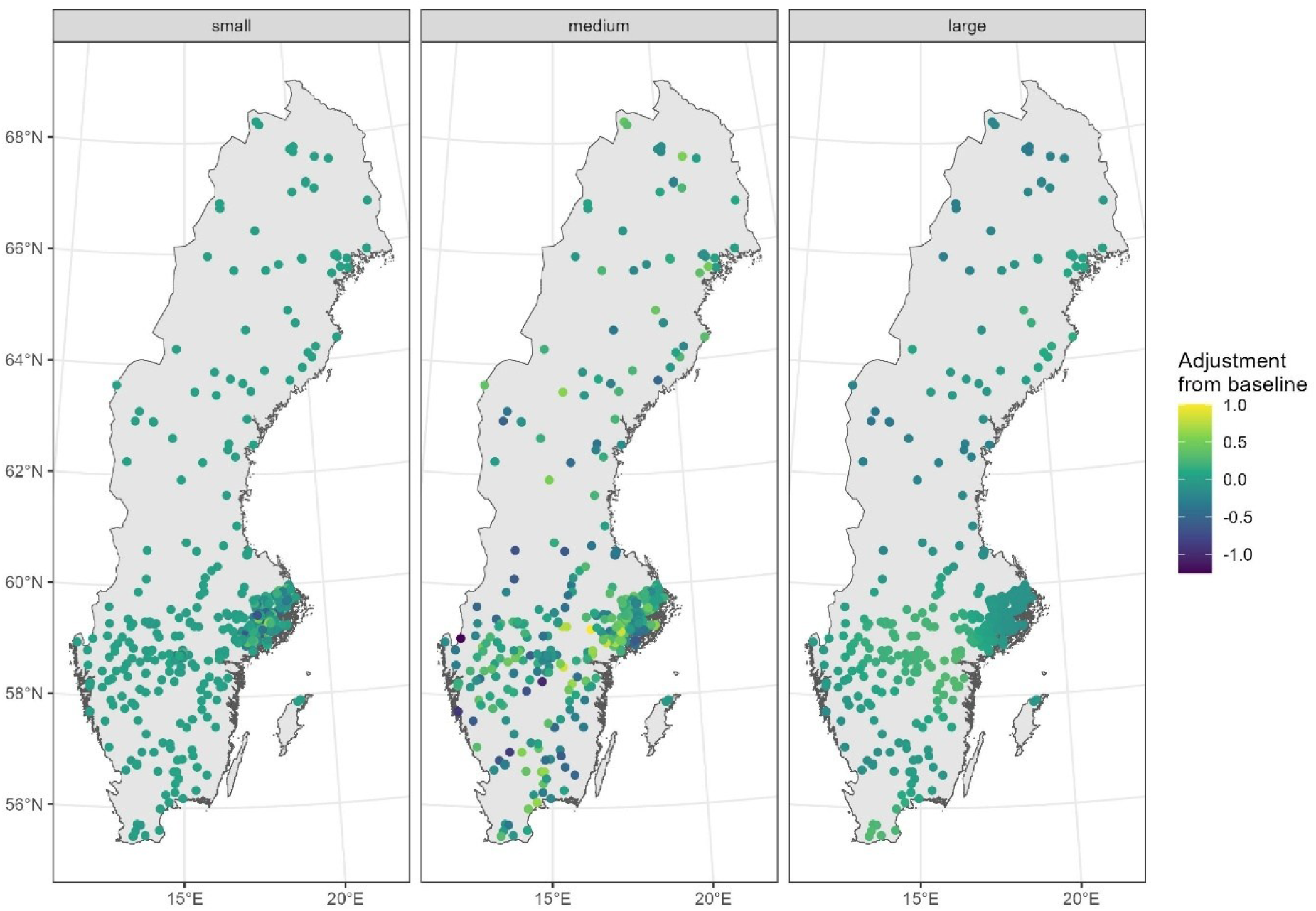
Null (no covariate, only random effect) model of species richness in all 425 Swedish lakes. CF-GLMM spatial components split into three equally-complex scales: small (2-10 km), medium (10–100 km), and large (100–460 km). Spatial scales adjust richness from its baseline, the non-spatial intercept β_0_, and adjustments are multiplied by bandwidth-specific distance weights before summing together into the predictive mean.

Generally, we got more precise predictions for sites with higher richness: these heteroscedastic patterns suggested underdispersion and limits to a classic Poisson approach (Figure 4). Temporally, we saw artefacts of the survey protocol with tighter fits in some intermediate years and fewer recent observations, especially after 2020.

**Figure 4:**
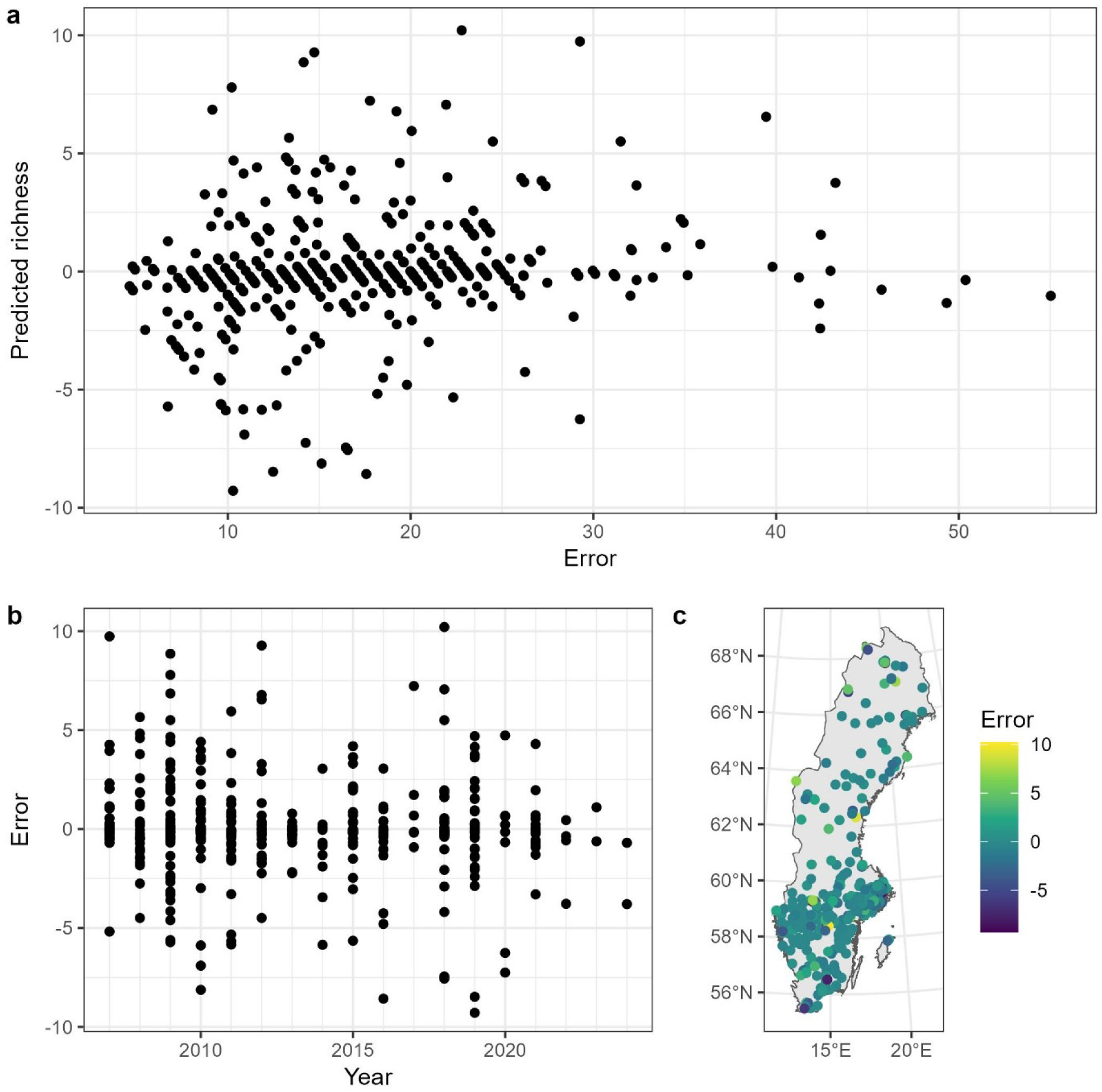
Goodness-of-fit exploration of the null spatial model for all 425 Swedish lakes without chemistry variables using raw residuals (observed - fitted values) to measure model error with a) general, b) temporal, and c) spatial trends. While spatially suitable, this model shows signs of general heteroscedasticity (i.e., underdispersion) and temporal trends, with the latter stemming in part from asymmetric surveying across timepoints, as fewer sites have their first surveys and more get resurveyed in later years.

Spatially, residual patterns distributed evenly across Sweden, suggesting that the CF-GLMM properly accounted for spatial autocorrelation in the data. Heteroscedasticity (i.e., error terms in our model do not have a constant variance across all observations) did not contribute to bias but may limit precision, error, and conclusions about significance. In short, inferential caution is warranted.

For the multiscale objective, we first look at the fixed effects. Models without squared terms appeared to operate on similar magnitudes — also referred to as “ecological significance” (Byrnes and Dee, 2025). TN appeared consistently positive, regardless of the additional terms included in the model, and its squared term neither affected other terms nor was significant itself (Figure 5a). TP seemed unstable overall, and its strong correlation with TN suggested that we should prefer TN to be cautious of collinearity (Figure S1). Temperature stood out — always statistically significant with changing directionality and larger magnitude when considered with a squared term (Figure 5a). For inferential purposes, a strong, nonlinear relationship between temperature and richness seemed a foregone conclusion: moderately higher temperatures increased richness while more extreme ones reduced it. Overall, the negative impacts dominated over the positive ones when simplifying to a linear effect.

**Figure 5:**
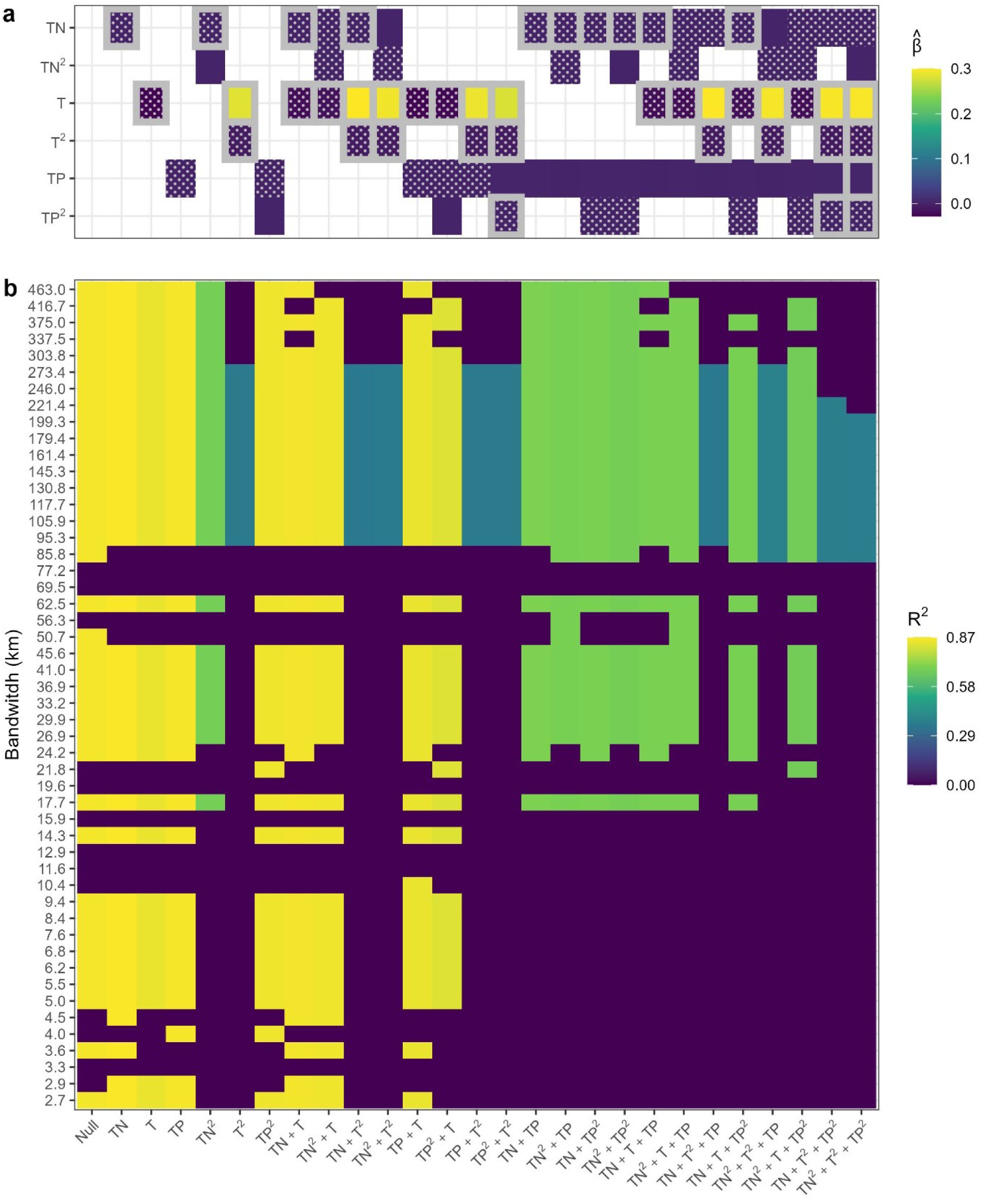
Model results plotted for 27 candidate models considering total nitrogen (TN), temperature (T), and total phosphorous (TP) along with their squared terms. a) Fixed effect coefficient mean estimates, with 95% statistically significant values surrounded by a grey box and negative values filled with a dot pattern. These indicate the expected percent increase in species richness from an additional 1°C or 1 µg of water temperature or TN/TP, respectively. b) Distance bandwidths identified as relevant to the model coloured with the model’s overall R^2^ value. A 0 value indicates that the bandwidth for that distance was omitted from the model. Squared covariates only appear when the base term is also in the model (e.g., a TN^2^ model has both TN and TN^2^ terms, and TN^2^ never appears without a complementary TN term).

In contrast, the predictive performance at various spatial scales emphasised different patterns than inferential interpretations. The filtered null model performed well, and most attempts at improvement yielded worse results (Figure 5b). The worst performance came from models with T^2^, and all trivariate models. A model with just TN proved most explanatory (0.009 R^2^ increase), followed by TP (0.003 R^2^ increase), but both offered only slight improvements over a null spatial-only model. Thus, we might consider if some missing component of geochemistry or human impact that affects nutrients was missing from the pure spatial effects. Still, we should be sceptical given only minor R^2^ increases despite the metric’s tendency to increase for more complex models.

Regarding the spatial confounding problem, we note that degraded explanatory performance arose after more complex models (i.e., including TN alongside either TN^2^ or TP), removing all small-scale spatial terms below 16 km. And T^2^ models further removed medium-scale bandwidth terms below 82 km. But focussing on high-performance models (i.e., > 0.8 R^2^), they generally covered comparable bandwidths on the small and medium scales, with some variation ±1 and ±10 km at the respective scales. Most interestingly, we saw that including both TN/TP and temperature in the same model always came at the expense of some large-scale bandwidths (i.e., 300+ km), supporting our interpretation that these large-scale processes represent larger macroecological gradients.

Overall, temperature and TN did not meaningfully predict species richness — the null model had the lowest average prediction error (i.e., ±0.1 species) and comparable R^2^ values (Table S1). These predictions hold despite variables’ inferential and ecological relevance. The missing predictive relationship follows from the baseline spatial patterns (Figure 2) and minimal improvements offered by additional variables to the flexible multiscale model (Figure 3).

## 4. Discussion

### 4.1. Data Utility

Spatial models and their inherent potential for spatial confounding can respond strongly to patterns in the data-generating process, and we note several possible areas of caution related to assumptions and accessibility (Khan and Berrett, 2024). First, our data cleaning assumptions (e.g., shallowest chemistry only) suit our exploratory research aims but should be cautiously applied for other research questions (e.g., depth-sensitive joint species distribution models); spatial scales and relevant confounding distances may change as a result.

For accessibility, digital systems often reflect arbitrary cutoffs due to sensitivity to data management decisions. In the Swedish case, the API portal only provides post-2007 WFD data despite the existence of earlier monitoring (Ecke, 2018). Other systems will have their own cutoffs alongside unique data structures requiring considerable time for developing replicable extraction pipelines. Documentation is often unavailable and language-limited when available, including in the Swedish case. Researchers may find similar data portals in their own countries that are functionally hidden from the research community due to linguistic inaccessibility and different open access protocols but present similar macro-scale research possibilities. Inconsistent definitions of macrophytes present another challenge that emerges when considering the difference between reported WFD richness and Murphy list richness . One approach may consider cyanobacteria and chlorophytes as macrophytes while excluding charophytes; another may apply the opposite standards, impeding consistent field sampling and whole-lake community representations (Chambers et al., 2008; Murphy et al., 2019).

Finally, sampling and survey methodology, both including and going beyond proxies for effort, can directly affect macrophyte richness estimates. These values, often missing from public databases, can complicate comparisons across jurisdictions (Pentecost et al., 2009). Relatedly, missing values present another persistent issue, and solutions can involve imputing, removing, or interpolating values (Larson et al., 2023). However, gap-filling can introduce bias in exchange for better precision, creating particular problems for inference (Haubrock et al., 2025). Instead, modern modelling approaches increasingly accommodate missing values, and using those can render such questions irrelevant if suitable for research aims (Korhonen et al., 2025; Skarstein et al., 2023). For our spCF multiscale analysis, we opted against filling missing values to avoid issues in interpretating fixed effects inferentially; if focussing just on prediction, the other interpretation would be more plausible.

### 4.2. Model Generalisability

Despite our concerns about model dispersion from residual inspection, we noted no difference between quasi-Poisson and Poisson distribution families. Since quasi-distributions keep the same β (i.e., fixed effect) estimates with different variances, spCF likely assigns this variation to the spatial random effect rather than dispersion term — a possible methodological limitation (Villadsen and Wulff, 2021). Overall, distributional limitations somewhat constrain models’ ability to fit ecological reality; for example, Tweedie (i.e., a family of probability distributions usually applied in ecology to data with many-zero, positive, and either discrete or continuous values such as species abundances) suits such count data well but is often incompatible with GLM-based models (Foster and Bravington, 2013).

Model fits also did not accept missing values, had scale sensitivity to offset terms, yet comfortably allowed non-normalised fixed effects (Appendix A). We attempted to incorporate binary fixed effects for each survey year (i.e., “econometric random effects” or “within cluster means”) to supplement limited CF-GLMM temporal flexibility but found that the resultant models selected no spatial trends — a known limitation where these terms absorb between-site gradient variation for smaller sample and cluster sizes and another illustration of pervasive confounding issues for inferential analysis (Byrnes and Dee, 2025). Such considerations arise especially for spatiotemporal processes and point to strong correlations among spatial and temporal structures: for example, temperatures increase across time but increase more strongly in mountain than lowland areas (Diggle, 2013). More complex models are needed to tackle this non-separability issue.

Regarding the counter-intuitive predictive results from squared terms, CF models aim to flexibly capture non-linear effects, including the exponential of fixed effects (Murakami et al., 2026b). Thus, the poor performance of TN and TP squared terms despite their ecological relevance adheres to the models’ expected performance. In contrast, high significance for squared temperature may stem from its unique ecological importance. But because of this importance, the model replaces many flexible CF components with a single, less nuanced fixed effect that degrades prediction.

### 4.3. Novelty and Benefits

Our pipeline provides one of the longest and broadest datasets on spatiotemporal macrophyte communities and their diversity in Sweden. It has numerous possibilities for (macro)ecological research, including evaluation of multiple physiochemical parameters. Robust in-situ data can support not only our simple decomposition but more advanced multiscale remote sensing assessments by helping to correct current models’ maxima underestimation bias (Putkiranta et al., 2025). Such pipelines will only grow more important as more countries embrace open access data policies, like Finland (http://tun.fi/HR.7080), the Netherlands (https://wkp.rws.nl/downloadmodule), and Japan (https://www.biodic.go.jp/moni1000/findings/data/index_file_LKaquaticplants.html). Despite our primary focus on and interest in macrophytes, Sweden’s Miljödata MVM portal also includes other WFD target taxa, including fish and macroinvertebrates, that can be extracted using our code with only slight modifications.

We demonstrate a workflow for WFD data entirely accessible and available via API — easy to access but more difficult to process. Other formats include static datasets (i.e., not updated with future years) and public portals requiring more complex scripting to extract usable data for modelling (Jonge et al., n.d.; Lewerentz et al., 2023). Live datasets can have restrictive terms of use (e.g., limits on raw data dissemination with unregistered users), so scripts provide a replicable path forward.

Analytically, our model provides key insights into the spatial confounding problem in linear models, where the spatial component impedes unbiased fixed effect estimates (Khan and Berrett, 2024). The spatial complexity of the non-parametric component can directly influence the magnitude of this issue even in complex models for both prediction and inference (Ward and Anderson, 2026). By showing a feasible implementation for direct estimation of relevant spatial scales and its responses to the progressive addition of fixed effect terms of interest, we demonstrate a mitigatory workflow for future spatial models to better tackle their underlying user-specified sensitivities, properties, and assumptions (Murakami et al., 2026a, 2026b).

Changing spatial scales between our single- and multiple-predictor models (i.e., TN in isolation or TN and T together) exemplify a complementary “spatial collinearity” paradigm (i.e., collinearity as one source of confounding). Here, the spatial interaction of two separate spatial terms can function on unique scales not represented in the processes alone — corresponding to a spatial-specific interpretation of limnological interactions like warming-eutrophication allied attack (Meerhoff et al., 2022). While we used a spatial random intercept to test spatial confounding between fixed effects and spatial terms here, spatial random slopes (i.e., spatially varying coefficient models) can circumvent the entire spatial confounding issue. However, they trade increased flexibility within a study area for worse generalisability outside of it (Doser et al., 2024).

Regarding suitability and interpretability, our model shows clear divergence in a “best” model for inference or prediction. Our null models accentuate the power of ecologically naïve approaches that exclude traditional chemistry variables entirely. Because of spatial confounding, including well-identified and significant variables can act against good prediction.

Nonetheless, both inference and prediction face challenges from spatial confounding. In inference, coefficients can have the wrong magnitude, direction, precision, or all of the above (Khan and Berrett, 2024). In prediction, modelling with the wrong spatial scale can drive underestimates of observation error, making a model overconfident and structurally degrading predictions for a particular data subset (e.g., a single species in a multi-species model with a distinct spatial behaviour or cluster) (Ward and Anderson, 2026).

Such challenges apply especially to freshwater macrophyte macroecology patterns: the mechanistic drivers of diversity do not equally distribute across space and exhibit strong, scale-dependent structures (Alahuhta et al., 2021b). In addition to the inherent spatial asymmetry of lotic systems that occupy only portions of inland landscapes, survey imbalances introduce additional issues for freshwater systems (Schaeffer et al., 2024). For robust models with confident, generalisable conclusions, we must identify functional spatial scales that underpin our estimates of key diversity processes and their drivers.

### 4.4. Applied Relevance

More generally for freshwater modelling, we show how to identify problematic spatial scales to ensure good separation between fixed and random effect terms through simultaneous and automatic (i.e., rather than individual and user-dependent) comparisons. Such approaches can function either as self-standing spatial analyses or sensitivity checks to ensure better specification of other relevant models (Murakami et al., 2026a, 2026b).

In WFD monitored Swedish lakes, ecologically realistic models (i.e., with squared terms) amplified the consequences of spatial confounding while naïve models (i.e., without squared terms) better utilised the multiscale flexibility of the CF models. This contrast relates directly to research objectives — ecological realism improves inference and worsens prediction, and naïve models work in the opposite. However, both approaches benefit from considering spatial confounding. CF-GLMMs identified distinct spatial scales for squared covariates: temperature operated below 90 km and above 300 km while TN and TP (i.e., trophic chemistry) operated only below 17 km.

We can interpret these practically or statistically. In the practical application, additional monitoring for Swedish nutrient chemistry may provide more value-added if focussing on lakes within 17 km of existing sites and interested in macrophyte-nutrient relationships. But if aiming to forecast future macrophyte richness, sampling may be more cost-effective if focussing only on macrophyte records without additional survey effort spent on chemistry variables. In the statistical application, operational scales will directly inform results (Khan and Berrett, 2024; Ward and Anderson, 2026). As an example, extending our analysis with a hierarchical SPDE model to account for sampling bias should avoid including both spatial terms denser than 17 km and trophic covariates. Such a model structure would suffer from spatial collinearity, thus artificially decreasing statistical and ecological significances for both types of parameters.

We demonstrate the unique value of the highly recent CF-GLMM approach to multiscale modelling for macrophyte ecology, and our work provides a practical guide to these models for not only our own objectives but also other applied and fundamental research questions (Table 2). CF-GLMMs can suit either prediction or inference but specifically apply to questions and data with a spatial component.

**Table 2:** Illustration of CF-GLMM applicability to various spatial ecological questions involving both inferential and predictive approaches. These models can be used either as the main analytical method or as an intermediate step to other methods.

| Question Type | Question | Multiscale Application |
| --- | --- | --- |
| Inference (associative) | Do submerged and emergent macrophytes show similar or different patterns across spatial scales (e.g., for dispersal)? | Fit species-specific null model CF-GLMMs and compare their relevant spatial bandwidths. |
| Inference (causal) | Do changing agriculture and forestry patterns drive macrophyte diversity when considering specific environmental gradients? | Fit a CF-GLMM with agriculture and forestry as fixed effects to identify how spatial bandwidths change relative to the null model. Then, fit a more complex spatio-temporal model (e.g., SPDE) using the CF-GLMM bandwidths to build the spatial structure. |
| Prediction | How are macrophyte communities expected to spatially change in future climate scenarios? | Fit a CF-GLMM with most relevant climate variables as fixed effects to model important community metrics. Re-run with various scenario forecasts of those climate variables to produce future forecasts. Further extend to temporal models using relevant CF-GLMM bandwidths for spatial structures. |
| Prediction | How well do long-term biomonitoring metrics generalise to other sites | Fit CF-GLMMs for the metrics of managerial interest. Ensure sufficient training and test data sets. Compare |
|  | and years that might be less regularly sampled? | test set RMSE (outlier-sensitive) and MAE (outlier-insensitive) scores to understand limitations of monitoring. Repeat while considering model ensembles and other machine learning approaches for higher bandwidth errors and instability. |

## 5. Next Steps

Gaps in macroecological research for macrophytes remain pervasive given data and expertise mismatches combined with a relatively smaller research community compared to other taxonomic groups (Alahuhta et al., 2021a). Our data can help address some of these gaps (e.g., understanding of elevational and latitudinal relationships) while the workflow itself can apply to preparing European data and other public, global data for analysis (Table 3).

**Table 3:** Summary of relevant gaps for spatial macrophyte analysis divided into monitoring and modelling components alongside recommendations to modernise workflows

| Method scope | Recommendation |
| --- | --- |
| Monitoring | Understand the current accessibility status of monitoring data |
|  | Compile and publishing data for public use (either internally or via external request by researchers), which may require processing for quality control and sensitive data exclusion (Larkin et al., 2026) |
|  | Provide accessible documentation and dedicated contacts for fielding queries, with multilingual options preferred |
|  | Explore interactive, public visualisations to encourage better management, research, and public engagement (Delaney and Larson, 2023; Riffat et al., 2023) |
| Modelling | Check for spatial confounding (i.e., spatial collinearity) in addition to normal collinearity — both influence robustness and results |
|  | Evaluate all model parameter spatial scales to better understand analytical relevance, especially when using aggregation-based models like GAMs and MEMs |
|  | Clarify research objectives (e.g., inference or prediction); then select and report models appropriately (Leek and Peng, 2015) |
| | For prediction, ensure robust model validation steps that limit overfitting to the test data. Compare models using appropriate metrics (e.g., $R^2$, RMSE, Accuracy) and not significant parameters |
|  | For inference, ensure that data and models adhere to the stricter assumptions (Correia et al., 2026). Match interpreted effect parameters to the proper spatial/temporal/inferential scale for generalisable conclusions |

While macroecology clearly demonstrates the issues with scale-sensitive processes, the impact of multiple spatial scales (e.g., point clusters within lakes) persist more broadly across observational ecological data (Khan and Berrett, 2024). At the same time, ecological modelling best practices have advanced substantially in recent years, opening up new opportunities for applications to macrophytes (Byrnes and Dee, 2025; Correia et al., 2026).

Here, we take the first steps to unearth an underexploited world of WFD data and its accessibility for global researchers from data accessibility to results. Unlike experimental data, these monitoring datasets must be processed, including assumptions that should be verified for suitability with field realities (Soga and Gaston, 2025). But if done properly, researchers will be increasingly able to answer questions about phenology, dispersal, human impact, climate, etc. on larger and more generalisable scales with a truly replicable pipeline. Portals like Sweden’s Miljödata MVM are regularly updated, enabling real-time updates, verification, and even live dashboards for public communication.

Macroecological analysis requires comparable data over multiple large-scale ecological gradients — conditions that can be challenging to meet with public data despite the breadth of monitoring. Macroecology also frequently considers the relevance of dynamic processes (e.g., climate, land use, seasonality) that can change both within and across years. Our data access pipeline directly connects verified data to potential analysists in a usable format, one of the first such applications in a European context.

Our multiscale spatial decomposition workflow, too, provides a versatile and interpretable way to consider the roles many spatial patterns and their ecological impacts. We quantify distances at which richness, nutrient chemistry, and temperature operate. Specific scales link to different processes: local scales to site and sampling gradient, and large scales to macroecological trends. By identifying the relevant scales to a research question, this workflow can support numerous analytical objectives.

While distances can guide management and analysis, interpretation is key. We first interpret our results inferentially, where we distinguish drivers well at the expense of substantial unexplained variation in richness. We then interpret predictively, where good fit overall comes at the expense of traditional predictors. To support their use, we provide illustrative guidelines to use and explain these models.

In general, answering scientific questions with data requires awareness of the inherent assumptions both in data processes and modelling approaches. Each model reflects core beliefs about how processes are generated, structured, and related. Monitoring data, which directly informs government compliance and management, may have real impacts on the natural world regardless of analytical naiveté. While they may be daunting, we must consider the added dimensionality and structures that layer on top when considering space and time (Diggle, 2013). Proper analysis of data, especially when collected over larger scales, starts by checking the spatial scale of patterns and their relationships — which we show are not always equivalent. These can then support traditional approaches with informed user specifications that hold true to real-world contexts.

## Appendix A: Glossary

**Inference**: understanding the direct relationship between 2 processes more generally, often with the intent of establishing causality

**Prediction:** drawing conclusions about unsampled observations, often generalising to conditions (e.g., in space and time) absent from the modelled data

**Confounding:** two processes in a model impede each other’s estimates **Spatial scale:** a specific resolution in space at which processes operate **Linear model:** a model for Y based on adding together some Xs

**Intercept:** describing the baseline value of Y (not depending on X) with some term β_0_

**Fixed effect:** describing X’s effect with just one number (term), an interpretable β value

**Smoothed effect:** describing X’s effect with many numbers (terms), where each term corresponds to some basis function that can be difficult to interpret alone

**Polynomial effect:** describing X process(es) using a few terms β for each polynomial degree (e.g., β_1_ for x_1_, β_2_ for x_1_^2^, etc) that can be interpreted as fixed effects – a special case of smoothed effects

**Random effect:** describing the unexplained randomness left over, often using a known structure (e.g., space, time, observational unit)

**Offset:** rescaling the entire model by some variable without any β (e.g., surveyed area as an offset will transform Y from species counts to species densities)

**Species richness:** simplest form of biodiversity, the number of species detected

**Residual (error)**: a common way of measuring “left over” model error for each observation using the differences between observed and fitted/modelled/predicted values

**Homoscedasticity**: a core assumption of GLMM-based models, where the residual error is constant for all observations

**Heteroscedasticity**: the opposite of homoskedasticity, where model errors show some structure (e.g., related to some spatial, temporal, or environmental gradient)

## 8. Author contribution statement

M.T.: data curation, formal analysis, methodology, project administration, software, visualisation, writing – original draft, conceptualisation, funding acquisition, and writing – review and editing

J.G.: supervision, conceptualisation, funding acquisition, and writing – review and editing

J.C.: software, conceptualisation, and writing – review and editing

X.J.: supervision, conceptualisation, funding acquisition, and writing – review and editing

D.J.L.: conceptualisation, funding acquisition, and writing – review and editing

J.A.: supervision, visualisation, project administration, conceptualisation, funding acquisition, and writing – review and editing

## Acknowledgements

We thank Andreas Rudh and Lars Sonesten for providing documentation and data clarifications in the Swedish WFD macrophyte and water quality sampling, respectively. Artificial Intelligence Generated Content (AIGC) tools were not used for writing, data analysis, or interpretation of findings. Finally, we extend our sincerest gratitude to the remaining project collaborators who could not be named in the present manuscript version.

## Data Availability Statement

All data are available through the Miljödata MVM data portal (https://miljodata.slu.se/MVM/) and subject to its terms and conditions of use. All can register an account with the portal to access an API token necessary for our scripts. Data import, processing, and analysis scripts are publicly available on GitHub (https://github.com/MishaTs/SwedenWFD). The repository also provides more detailed information on account registration, data access, and workflow.

## Supplementary Information

**Figure S1:**
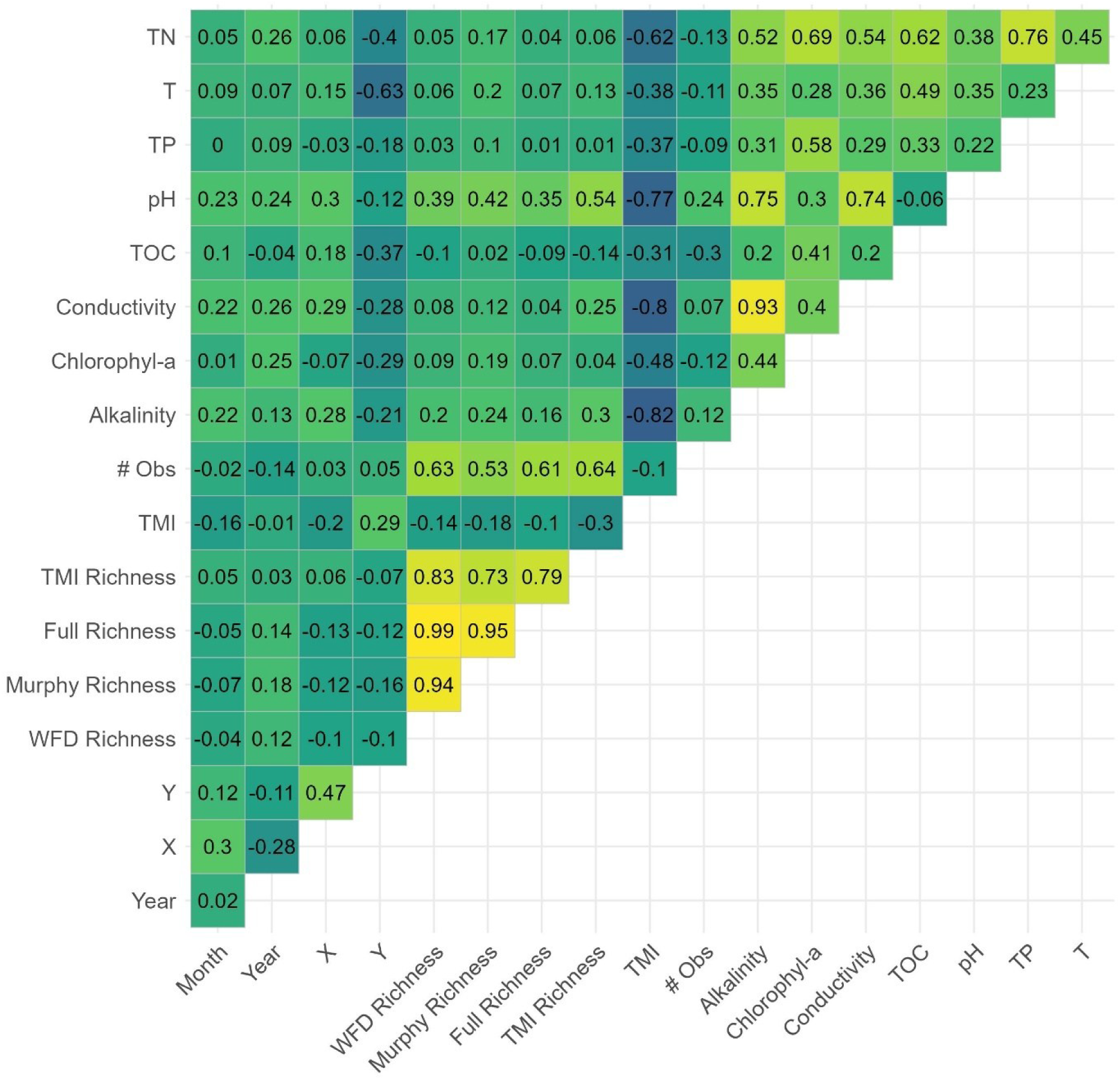
Correlation of main quantitative variables included in the final data, including sampling coverage, richness types, and chemistry parameters. X is the east-west coordinate, and Y is the north-south coordinate in the original SWEREF99 CRS (EPSG: 3009). See Table 1 for description of variable units and summary statistics.

**Table S1:** All available predictive scores for each model – pseudo R^2^, RMSE, and MAE. Pseudo (McFadden’s deviance-based) R^2^ ranges between 0 and 1; larger values are better. RMSE is overly sensitive to outliers and unitless; smaller values are better. MAE may underweight outliers but is in units of Y (number of species observed at a site); smaller values are better. Each metric’s smallest value is bolded and italicised, which matches the null model for MAE and the TN model for R^2^ and RMSE. See Figure 5 caption for more thorough model descriptions.

| Model | Pseudo $R^2$ | RMSE | MAE |
| --- | --- | --- | --- |
| Null | 0.8564621 | 3.154810 | ***0.1035073*** |
| TN | ***0.8658943*** | ***2.996811*** | 0.3365266 |
| T | 0.8436191 | 3.258931 | 0.1830217 |
| TP | 0.8597485 | 3.117811 | 0.2017968 |
| TN<sup>2</sup> | 0.6820693 | 4.653194 | 0.5838087 |
| T<sup>2</sup> | 0.3605152 | 6.778128 | 0.4880725 |
| TP<sup>2</sup> | 0.8487776 | 3.246039 | 0.2106707 |
| TN + T | 0.8556473 | 3.078039 | 0.3404916 |
| TN<sup>2</sup> + T | 0.8492738 | 3.175071 | 0.3900474 |
| TN + T<sup>2</sup> | 0.3506075 | 6.838875 | 0.4691967 |
| TN<sup>2</sup> + T<sup>2</sup> | 0.3577008 | 6.794620 | 0.5134554 |
| TP + T | 0.8424139 | 3.246385 | 0.2361359 |
| TP<sup>2</sup> + T | 0.8240647 | 3.419916 | 0.1612824 |
| TP + T<sup>2</sup> | 0.3599254 | 6.792308 | 0.4770797 |
| TP<sup>2</sup> + T<sup>2</sup> | 0.3588012 | 6.777395 | 0.5880085 |
| TN + TP | 0.6993865 | 4.464523 | 0.5482009 |
| TN<sup>2</sup> + TP | 0.6909742 | 4.550000 | 0.5338960 |
| TN + TP<sup>2</sup> | 0.6946706 | 4.515525 | 0.5626803 |
| TN<sup>2</sup> + TP<sup>2</sup> | 0.6814149 | 4.642200 | 0.5741292 |
| TN + T + TP | 0.6955562 | 4.484081 | 0.5124845 |
| TN<sup>2</sup> + T + TP | 0.6948455 | 4.492972 | 0.5095254 |
| TN + T<sup>2</sup> + TP | 0.3503748 | 6.839571 | 0.4716855 |
| TN + T + TP<sup>2</sup> | 0.6911671 | 4.530875 | 0.5333174 |
| TN<sup>2</sup> + T<sup>2</sup> + TP | 0.3816636 | 6.683544 | 0.5402420 |
| TN<sup>2</sup> + T + TP<sup>2</sup> | 0.6772340 | 4.644141 | 0.5871027 |
| TN + T<sup>2</sup> + TP<sup>2</sup> | 0.3741876 | 6.723644 | 0.5785667 |
| TN<sup>2</sup> + T<sup>2</sup> + TP<sup>2</sup> | 0.3708736 | 6.748296 | 0.5579399 |

